# Validating the representation of distance between infarct diseases using Word2Vec word embedding

**DOI:** 10.1101/2022.06.21.496958

**Authors:** Daiki Yokokawa, Kazutaka Noda, Yasutaka Yanagita, Takanori Uehara, Yoshiyuki Ohira, Kiyoshi Shikino, Tomoko Tsukamoto, Masatomi Ikusaka

## Abstract

**Objective:** To determine if inter-disease distances between word embedding vectors using the picot-and-cluster strategy (PCS) are a valid quantitative representation of similar disease groups in a limited domain.

**Materials and Methods:** Abstracts were extracted from the Ichushi-Web database and subjected to morphological analysis and training using the Word2Vec. From this, word embedding vectors were obtained. For words including “infarction”, we calculated the cophenetic correlation coefficient (CCC) as an internal validity measure and the adjusted rand index (ARI), normalized mutual information (NMI), and adjusted mutual information (AMI) with ICD-10 codes as the external validity measures. This was performed for each combination of metric and hierarchical clustering method.

**Results:** Seventy-one words included “infarction”, of which 38 diseases matched the ICD-10 standard with the appearance of 21 unique ICD-10 codes. The CCC was most significant at 0.8690 (metric and method: euclidean and centroid), while the AMI was maximal at 0.4109 (metric and method: cosine and correlation, and average and weighted). The NMI and ARI were maximal at 0.8463 and 0.3593, respectively (metric and method: cosine and complete).

**Discussion:** The metric and method that maximized the internal validity measure were different from those that maximized the external validity measures; both produced different results. The Cosine distance should be used when considering ICD-10, and the Euclidean distance when considering the frequency of word occurrence.

**Conclusion:** The distributed representation, when trained by Word2Vec on the “infarction” domain from a Japanese academic corpus, provides an objective inter-disease distance used in PCS.

## 1 Introduction

A common clinical reasoning strategy is to recall a disease and check whether the patient history obtained is consistent with the disease.[1] Physicians take the first step, i.e., disease recall, in terms of prior probabilities that vary according to age and sex and the function and location of the medical facility. Based on these prior probabilities, they often recall one or two diagnoses and consider them based on a patient’s symptoms and characteristic findings in clinical practice. Sometimes, a list of differential diagnoses is required as competing hypotheses for a given diagnosis. Differential diagnosis lists are generated by specific rules, such as diseases with other pathologies occurring in the same organ, diseases occurring in anatomically adjacent organs, and diseases with similar pathologies but occurring in multiple organs. However, when physicians generate disease recall and differential diagnosis lists, there is always a chance for diagnostic errors due to heuristic bias.

The pivot and cluster strategy (PCS) can be used to avoid such bias,[2] in which clinicians simultaneously recall a differential diagnosis list that approximates one of the recalled diagnoses based on intuition. The process involved in PCS is as follows.[2, 3] First, a clinician designates the initial diagnosis (pivot) as the most likely hypothesis through an intuitive or analytical process based on history and physical examination, their knowledge, and experience. Second, the clinician forms a disease cluster around the pivot to obtain a collection of differential diagnoses. Any disease can be a pivot. The list of differential diagnoses can be reduced or expanded according to the overlaps and differences between pivots and their clusters.

Pivot designation is intuitive and can lead to diagnostic errors due to heuristic cognitive bias. The frequency of disease, which depends on the function and location of the medical institution, and the physician’s specialty and case experience influence pivot designation.[4] However, Shimizu et al. argued that in PCS, the automatic and simultaneous recall of clusters close to the pivot’s clinical presentation removes bias and improves diagnostic accuracy by preventing early closure.[2] They stated that PCS is also useful when teaching novice students. In PCS, the teacher, an experienced clinician, assigns a virtual distance from the pivot to the differential diagnosis and translates the list of differential diagnoses into a two-dimensional visual representation. This helps the learner represent the degree of concurrence with the patient’s clinical symptoms.

Shimizu et al. stated that, in PCS, pre-prepared clusters can automatically and quickly be recalled in batches. Nevertheless, a question arises as to “what do they mean by ‘pre-prepared clusters’ in actual clinical practice?” We contend that it may be based on the accumulated intuition that each clinician has experienced or the influence of textbook knowledge. In general, disease clusters based on similar pathophysiologies are likely to have similar medical characteristics. Despite physicians being empirically aware of “similarities” among disease groups, there has been no quantitative presentation of these disease groups. Exhaustive clusters are required because user-friendly, limited clusters based on physicians’ experience can be a source of cognitive bias. However, it is difficult to prepare uniform clusters for all pivots, and even if one could, it would be impractical to use them for a physician’s reasoning. Conversely, the differential diagnosis generator is a computer-generated list of differential diagnoses, which reportedly allows clinicians to reconsider their diagnoses.[5] If we can quantitatively represent clusters, the accuracy of the differential diagnosis generator is expected to improve significantly.

We collected documents related to “infarction” from the corpus of Ichushi Web, a database of medical articles, used Word2Vec to learn word associations from the collected articles, and found that the pathophysiological and anatomical features of “infarction” are retained in the distributed representation.[6] In our previous article (under peer review), it is suggested that “brain infarction” and “myocardial infarction” have different vectors and some vectors share greater similarity if they are diseases of the same organ. Word2Vec is an unsupervised learning system that uses neural networks and a tool to compute distributed representations of words.[7] Word2Vec is effective in capturing semantic relatedness and similarity relations among medical terms.[8]

This study examined the validity of the similarity (inter-diseases distance) calculated using our learned word embedding vectors as candidates for clusters in PCS. To evaluate the validity, we verified the agreement rate with the International Statistical Classification of Diseases and Related Health Problems (ICD), an external classification list provided by the World Health Organization. The 10th edition (ICD-10) is available in Japanese. It contains codes for diseases, their signs, and symptoms. Its hierarchical structure is divided into 22 chapters, and the circulatory system diseases (I00-99) include many strokes and myocardial infarctions. It is highly reliable because it is based on human judgment informed by pathology and anatomical relationships.

Would clustering by word embedding vectors be similar to ICD-10 clustering? If demonstrated, this hypothesis would provide reasonable grounds for a quantitative and objective presentation of clusters by word embedding vectors in medical corpora.

The remainder of this paper is organized as follows. Section 2 discusses the corpus and training methods used to obtain the word embedding vectors and the method of evaluating their validity. Section 3 presents the results of this study. Section 4 discusses the differences in distance definitions and update methods that maximize each validation scale, with reference to the dendrograms, the applicability of this study to clinical medicine, and the study limitations. Finally, Section 5 concludes the study.

## 2 Materials and methods

### 2.1 Subjects

We extracted abstract texts from the medical journal database of the NPO Japan Medical Abstracts Society, Ichushi (it requires registration), by searching for the word “infarction,” and we set the search criteria to show case report articles that contain an abstract.[6] We limited the infarction domain because it involves the same pathological changes in multiple organs. Many researchers have used abstracts from Ichushi web as subjects for their literature review.[9, 10, 11] We inserted spaces between words and performed morphological analysis on the abstract texts to produce word sequences converted into standard form. In Japanese, unlike English, there is no space between words. Therefore, to separate words, it is necessary to insert a space while checking against a dictionary. MeCab was used for inserting the spaces and morphological analysis, and mecab-ipadic-NEologd[12] and ComeJisyo[13] were used for the dictionaries.[14] ComeJisyo is a dictionary for MeCab that leaves a space between the words that represent terms used in medical facilities. After morphological analysis, we exclusively extracted terms consisting of nouns, adjectives, adverbs, and verbs. Numeric expressions are typically not crucial in Japanese natural language processing. We excluded a number of nouns from the sequences, e.g., numerals indicating a subject’s height and weight, and the dosage of medication.

### 2.2 Model learning

We trained the word sequences using the Gensim Word2Vec package.[15] Word2Vec is available in two different forms: skip-gram and continuous bag of words (CBOW). In previous reports, Word2Vec, specifically the skip-gram architecture, achieved the highest score on three of four rated tasks; analogy-based operations, odd one similarity, and human validation.[16]. Skip-grams also performed better in biomedical studies.[17, 18, 19, 20]

This study used a skip-gram algorithm based on previous studies, with a vector of 200 dimensions. We set the window size as 5, minimum count as 5, and the number of iterations as 100. Words containing “infarction” were extracted from the vocabulary used for training. We kept only the words found in ICD10 that matched perfectly and used the set of words as the standard name list.

### 2.3 Clustering and evaluating with internal validity measures

Each word in the standard name list had a vector of 200 dimensions. We used the hierarchical clustering in ICD-10 based on each vector component. As the figures in later sections show, (the terms in parentheses follow a lower-case naming convention) we used distance definitions (metric), including Euclidean (euclidean), city-block (cityblock), standard Euclidean (seuclidean), cosine, correlation, Chebyshev (chebyshev), Canberra (canberra), and Brady Curtis (braycurtis) distances. We used update methods (method), including single-linkage (single), complete linkage (complete), group average (average), weighted average (weighted), median, Ward’s (ward), and centroid. With this combination of metrics and methods, we first calculated the cophenetic correlation coefficient (CCC), which is an internal validity measure of clustering. We then created a dendrogram using the combination with the highest CCC value.

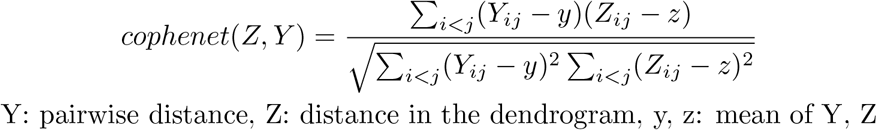

### 2.4 Evaluation with external validity scale

We evaluated the model with an external validity scale. External validity measures performance by matching the clustering structure to a priori information, i.e., true class labels.[21, 22] Each word in the standard name list was assigned its corresponding ICD-10 code. We counted the number of unique ICD10 codes and calculated the adjusted rand index (ARI),[23, 24] normalized mutual information (NMI),[25] and adjusted mutual information (AMI)[26] as external validity measures when clustering the standard name list to that unique number. These are the most commonly used evaluation measures to assess the similarity between two sets.[27] The range of ARI is [-1,1], that of NMI is [0,1], and that of AMI is [0,1]; a larger value indicates a closer match between the two groups. AMI is appropriate when the clusters are of unequal sizes and contain smaller clusters than ARI.[28] We calculated the distances and external validity measures for each combination while the update methods are those used in 2.3.

### 2.5 Analytics

The analysis was performed on a terminal using Ubuntu 20.04.2 LTS, with an Intel Core i9-9960X CPU, and 64 GB of primary memory; an NVIDIA GPU with 48 GB RAM, two TITAN RTX graphics accelerators, and an NVlink-bridge were used for computation. The machine learning framework was Python (3.6.9), Gensim (3.8.3), and scikit-learn (0.24.2), which is a module for machine learning in Python.

## 3 Results

Since October 26, 2019, 15,513 abstracts have been extracted (Ichushi ID: 1983011395 to 2019316513). Leaving space (as explained in subsection 2.1) and morphological analysis lead to 1,505,041 words and 46,602 unique words in the unlearned word sequences. The word sets containing “infarction” had 20,918 words, of which 71 were unique. The word sequences used for training Word2Vec had 1,445,433 words, with 15,163 being unique. The word sets containing “infarction” and used for training Word2Vec had 20,877 words, of which 54 were unique.

A summary of the set of words containing “infarction” extracted exclusively from words that fully matched ICD-10 (the standard name list), the total number of words, and ICD10 codes are shown in Table 1. The total number of words was 15,163, and unique words were 38. Twenty-one ICD-10 codes appeared; 16 of the 54 did not match the ICD-10 code. The words containing “infarction” that did not match ICD-10 were “infarction,” “hemorrhagic infarction,” “inferior infarction,” “post-myocardial infarction,” “postinfarction,” “inferior myocardial infarction,” “anterior infarction,” “anterior myocardial infarction,” “right ventricular infarction,” “infarctive,” “reinfarction,” “anterior septal myocardial infarction,” ““pulmonary infarction disease,” “impending myocardial infarction,” “multi-infarct dementia,” “perioperative myocardial infarction,” “thrombotic infarction,” “impending infarction,” and “posterior myocardial infarction.”

**Table1.**
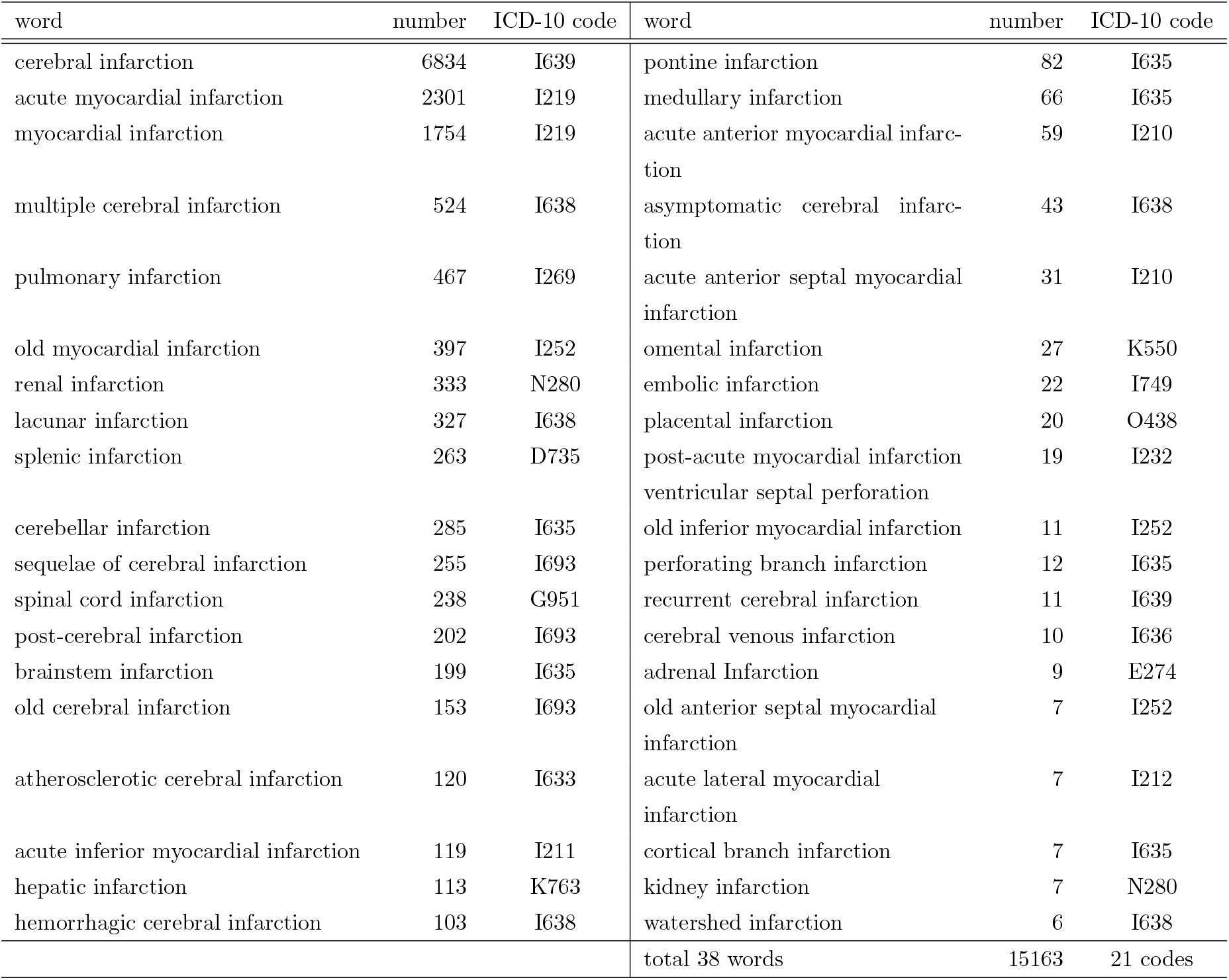
Standard name list: set of words containing “infarction” that precisely match ICD-10

Figure 1 shows the CCCs for each definition of the metric and update methods for the evaluation using the internal validity measure. When the metric was euclidean and the method was centroid, the CCC was maximal at 0.8690.

**Figure1.**
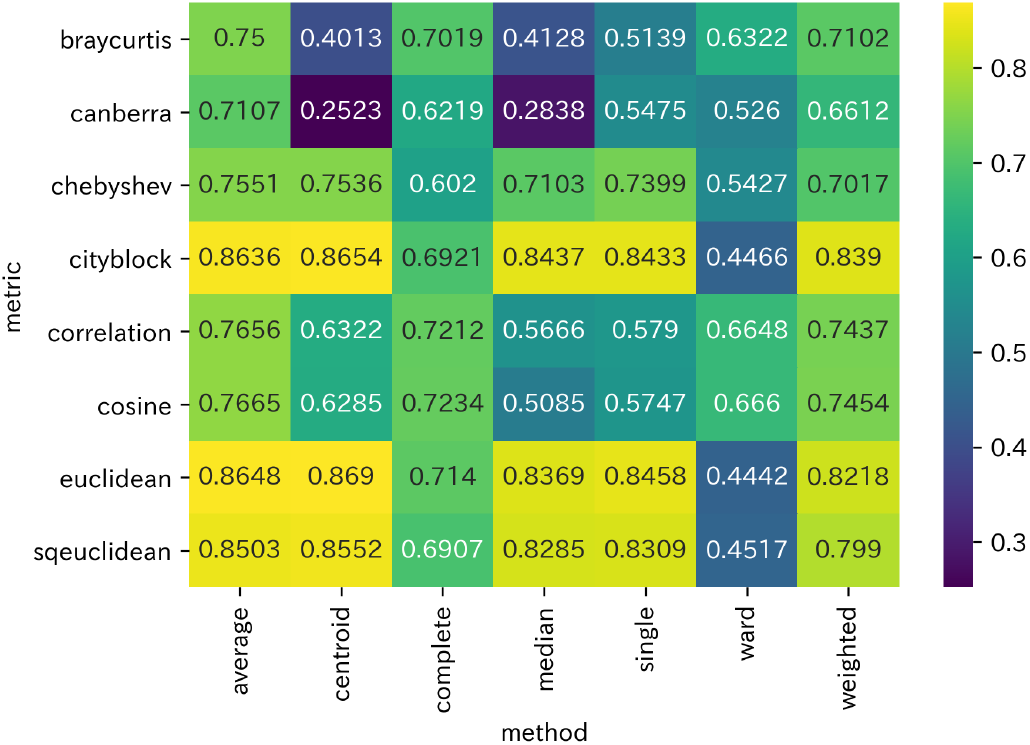
Internal validity measure with cophenetic correlation coefficients)

The evaluation using the external validity scale is shown in Figure 2. The AMI was maximal at 0.4109 with the cosine or correlation metric and the average or weighted method. The NMI and ARI were maximal at 0.8463 and 0.3593, respectively, with the cosine metric and the complete method. Combinations with high ratings on the internal validity scale did not necessarily have high ratings on the external validity scale.

**Figure2.**
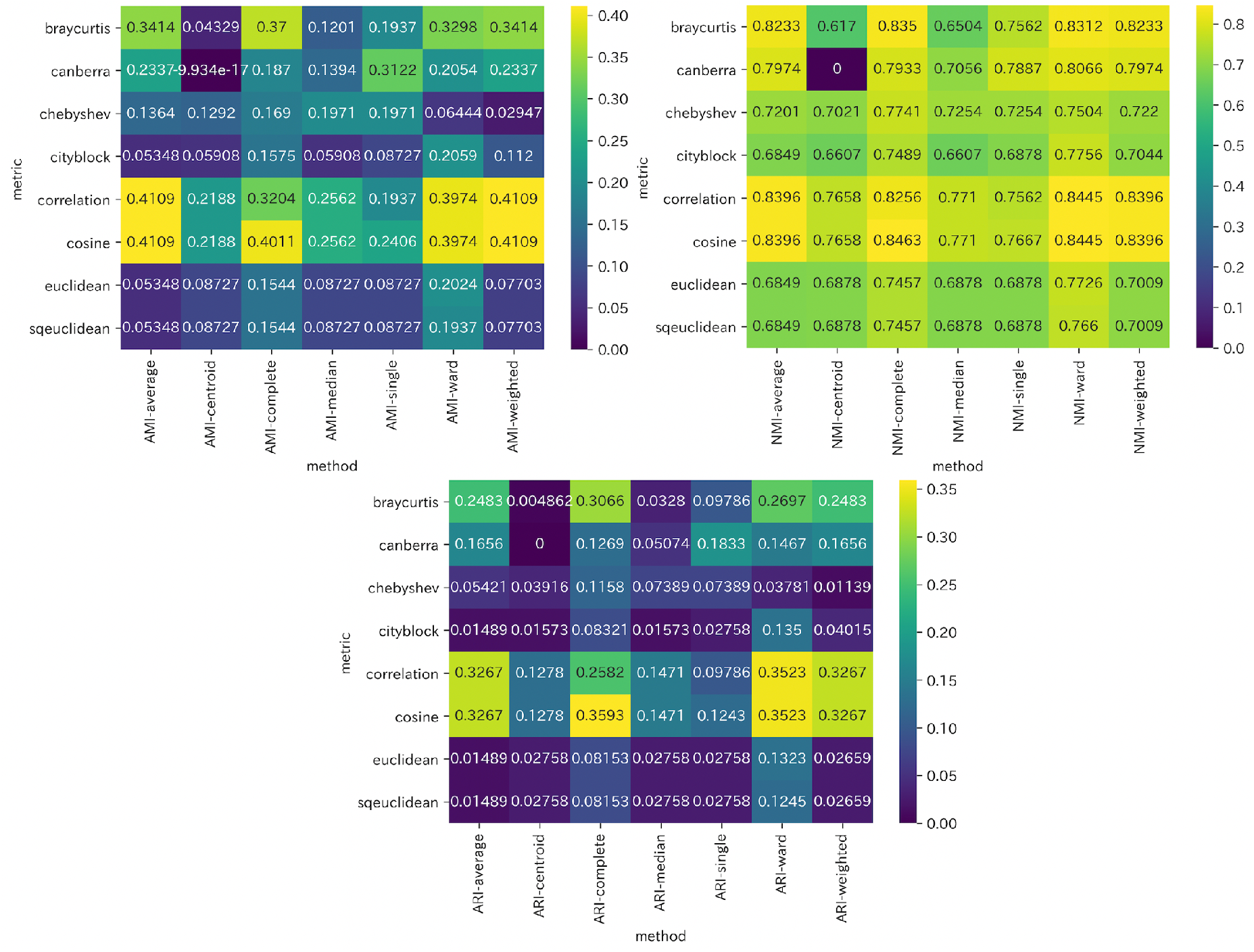
External validity scale for ICD-10

The dendrograms in Figures 3 and 4, discussed in detail below, illustrate the combination of metrics and methods that performed best on the internal and validity scale evaluations. In Figure 4, we arbitrarily set thresholds and colors to make the figure easier to view.

**Figure3.**
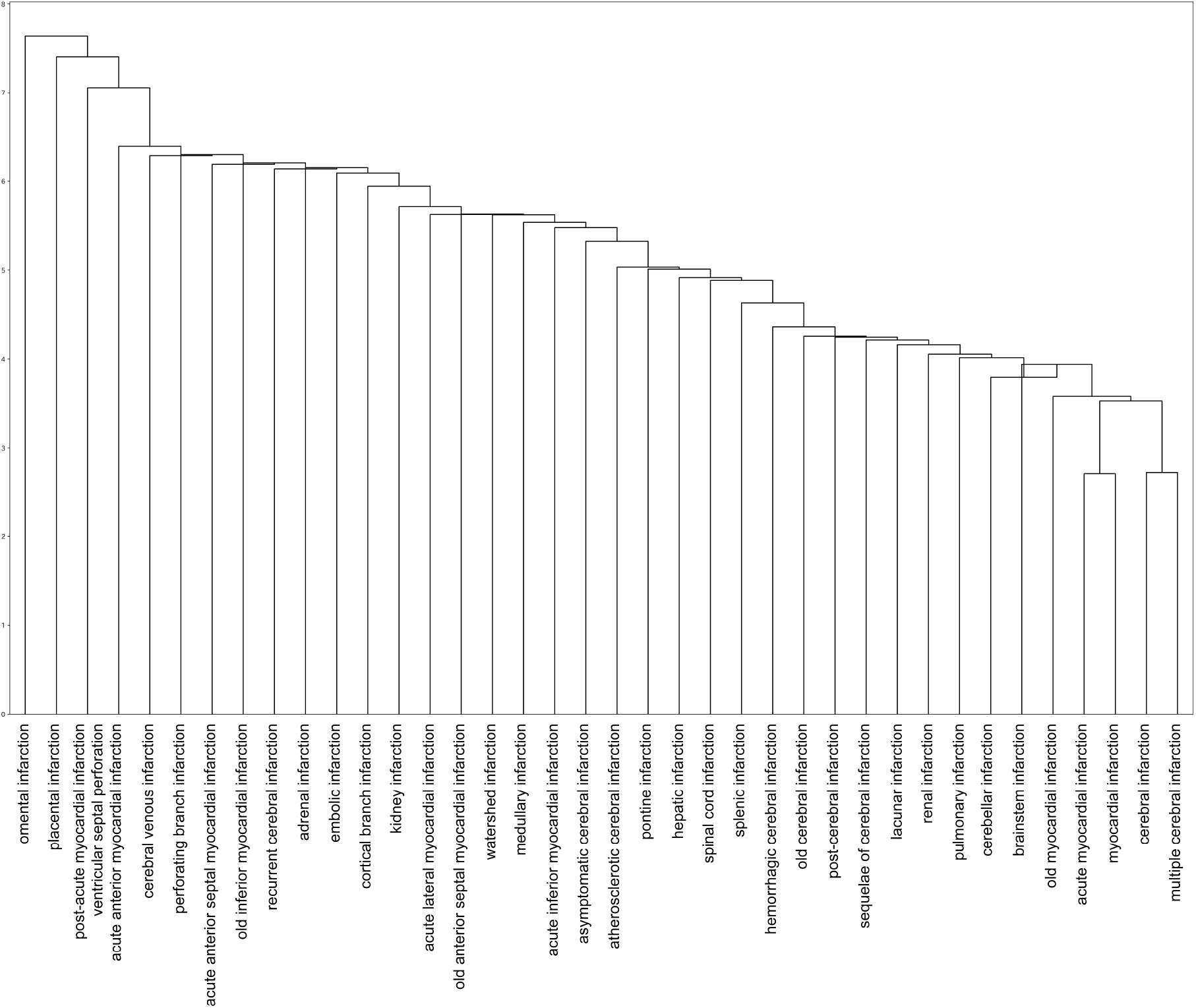
Dendrogram with euclidean and centroid that maximizes the internal validity measure. The vertical axis represents distance.

**Figure4.**
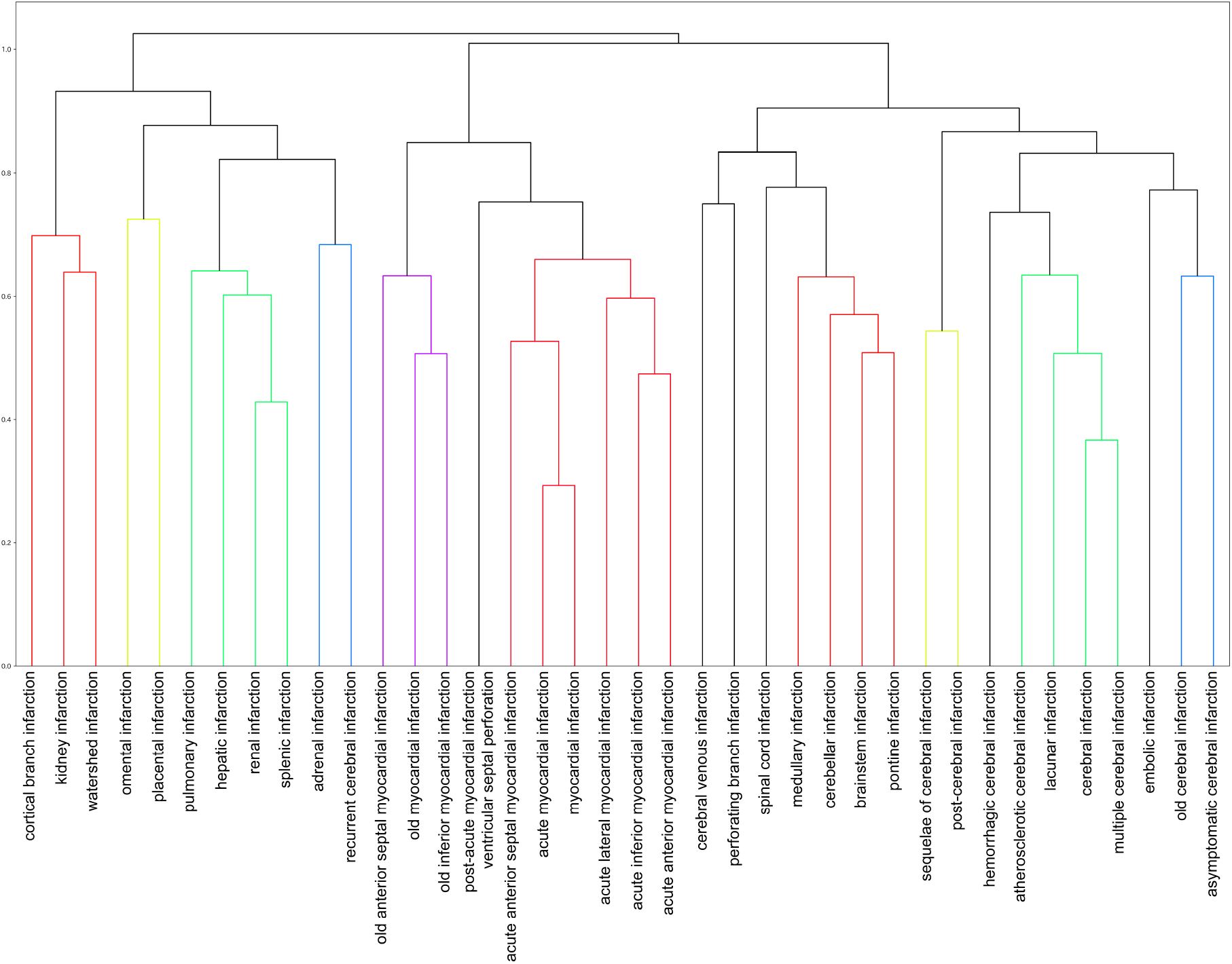
Six colored dendrogram when cosine and complete maximize the NMI and ARI. The vertical axis denotes distance, with 0.73 as the threshold.

## 4 Discussion

We extracted abstracts related to “infarction” from a database of Japanese medical documents and used them as a corpus to obtain word variance representations using Word2Vec. The variance representation thus obtained allowed us to measure inter-disease distances, which indicate the degrees of similarity among diseases. Our examination of multiple metrics and methods revealed that the combination of the euclidean metric and the centroid method was optimal for assessing internal validity, while the combination of cosine distance and the complete linkage method was optimal for assessing external validity with ICD-10 for NMI and AMI. The inter-disease distances between word embedding vectors is, therefore, expected to be a valid quantitative representation of similar disease groups.

Word2Vec uses deep learning based on the co-occurrence of words within a context to obtain a word embedding vector. Thus, words that appear in similar contexts will have high similarity. In academic abstracts, the description of infraction in an organ also includes clinical symptoms and characteristic information derived from that organ. Differences in the characteristic information co-occurring in different organs may be a factor in the distance between diseases.

In the medical field, there have been few challenges to classification tasks with embedded representations. In a study that used a distributed representation obtained from medical records as visit embedding, the k-means method was used to classify the characteristics of patients by specialty.[29] A German Word2Vec model trained on a corpus of 352 megabytes of medical reports attained an accuracy of 90% in assigning medical reports written by physicians to ICD-10.[30] The study identified rare diseases, unusual designations, and ICD code degeneracy as sources of assignment or “missing” errors. ICD-10 has a hierarchical structure with more than 68,000 codes. However, not all codes have the same level of granularity. Furthermore, it is known that ICD codes that are ontologically distant are less likely to be grouped.[27] Although we have bound our embedding vectors to ICD-10 based on the corpus of academic literature, the maximum value of NMI is 0.85 but only 0.41 and 0.36 for AMI and ARI; therefore, our embedding vectors cannot be interpreted as a mapping to the continuous space of ICD-10.

It should be noted that the metric and method that maximized the internal validity measure and those that maximized the external validity measure produced different results. The euclidean maximized the internal validity measure but did not maximize the external validity rating. The dendrogram with the parameter that maximizes internal validity (Figure3) gives the impression that, unlike the ICD-10 classification, the less clinically relevant diseases are adjacent. Conversely, the dendrogram with parameters that maximize the NMI and ARI with ICD-10 (Figure4) gives the impression of a classification based on anatomical differences and temporal differences such as “old” and “sequelae.” In other words, the latter could be classified as “ICD-10-like.” In other words, one would feel that the latter could be classified as ICD-10-like.

The Euclidean distance is calculated between two points as

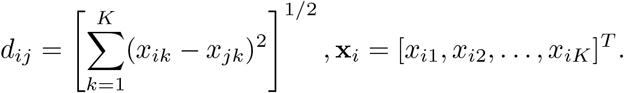

Conversely, the cosine distance is calculated from the angles between the vectors.

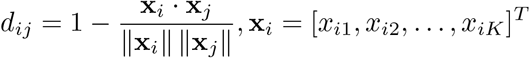

When performing calculation with cosine distance, the length information of the vectors is lost because it is divided by the L2 norm. In other words, when the angle between two vectors θ is 0°, the cosine distance is 0 (similarity is 1) even if the L2 norm is different; therefore, diseases at different distances from the origin (L2 norm) in a higher dimensional space are expressed as being similar. The L2 norm of **x**_*i*_ is calculated by the formula

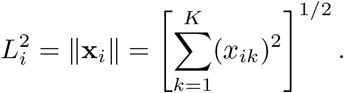

When clustering by cosine distance, all disease embedding vectors are normalized and plotted on the n-dimensional hypersphere, losing the L2 norm. Pivot and cluster were converted into a two-dimensional representation and shared with others, but this could also be represented on the hypersphere. A cluster is a group of diseases whose angle with the pivot is *θ* < *θ*_0_, and the inter-disease distance can be defined as the distance between two points on the hypersphere.

Conversely, the L2 norm is not negligible. Vectors represent words that are consistently used in similar contexts with larger L2 norms than words of the same frequency used in different contexts.[31] Applied to diseases and symptoms, medical words that denote a verity of causes, are highly abstract, and are used in various contexts (e.g., “infarction” and “headache”) have a smaller L2 norm. By contrast, medical words that are less diverse in terms of the cause, are more specific, and are used in limited contexts (e.g., “omental infarction” and “placental infarction”) have a more significant L2 norm. In fact, the L2 norms of the embedded vectors in this study are “infarction”:3.107, “headache”:3.955, “omental infarction”:7.703, and “placental infarction”:7.583. The L2 norm of the standard disease list is shown in Figure 5. The frequency of occurrence and the L2 norm tend to be inversely proportional. The similarity with the dendrogram shown in Figure 3 is also clear. Because the L2 norm is a Euclidean distance, this dendrogram reflects the frequency of occurrence and context of words in the corpus. Thus, using an unnormalized vector for classification may be preferable for considering word frequency and polysemy and to apply PCS in actual clinical practice.

**Figure5.**
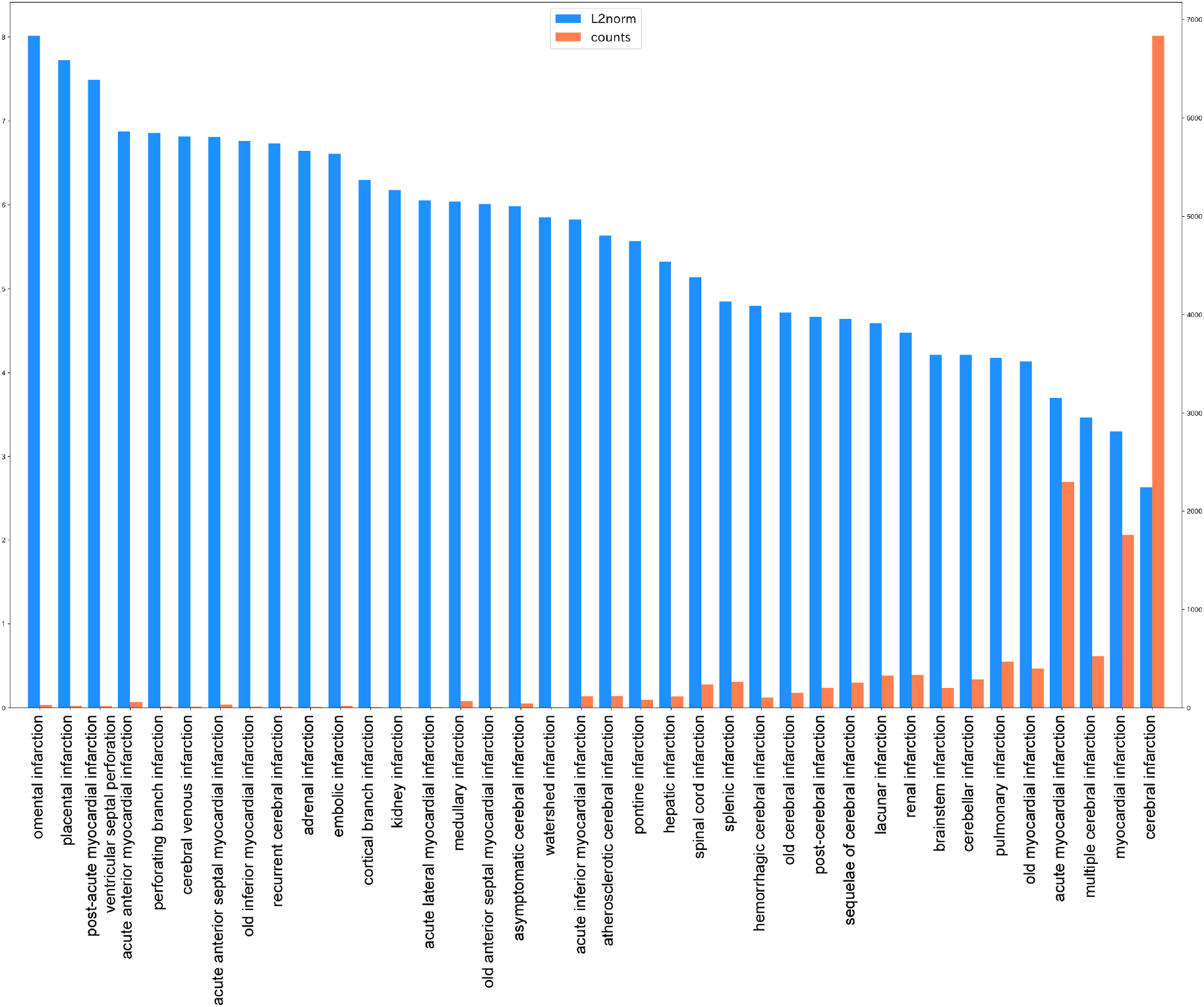
L2 norm and word counts of embedding vectors

The word frequencies should be limited to and interpreted within the corpus used in this study. However, using a broad corpus of academic medical literature may, in principle, be consistent with disease frequencies in the real world. Prior probabilities are essential information in clinical reasoning. By using Bayes’ theorem to modify the posterior probability of a diagnosis when new information becomes available, the prior probability represents its starting probability. In many cases, the prior probability depends on the function and location of the medical facility. By leaving the L2 norm to obtain clusters, a differential diagnoses list that considers prior probabilities in the corpus or facility may be obtained. By using different clusters for different stages of clinical reasoning (varying the distance and updating method), such computation may provide a more efficient differential diagnoses list that is more in line with the physician’s thought process.

Although this study was limited to the infarction domain, depending on the dictionary used in the morphological analysis, we simultaneously obtained the embedded vectors of medical words other than infarction disease. In other words, we efficiently computed vectors that represent symptoms, such as hemiplegia and headache, and histories such as smoking and hypertension. In the future, we will examine validity scales for domains other than infarction and calculate inter-symptom distances or symptom-disease distances to visualize many keywords used in clinical reasoning.

### 4.1 Limitations

There are several limitations to this study. First, we cannot assert that the corpus size used for the study was sufficiently large. Previous studies using academic literature corpora have acquired over 18 million abstracts to obtain a vocabulary of approximately 7.8 million words.[32] This study found that increasing the size of the dataset does not necessarily improve the performance in measuring similarity and relevance. However, the accuracy increased proportionally with size up to a point. Because this study used all of the searchable medical journals on Ichushi web, a different resource should be considered to increase the corpus size. Second, morphological analysis presents a problem. Many medical terms consist of multiple words, which is also true in Japanese; for example, “acute inferior myocardial infarction” contains four words in English and eight Chinese characters but is a single medical term in both languages. Word2Vec is vulnerable when encountering unknown words, and if a term is not entered as a multi-word term in the dictionary, it is divided similar to the longest words. For example, if the medical term “acute right renal infarction” is present in the corpus but not in the dictionary, it will be divided into “acute,” “right,” and “renal infarction.” For morphological analysis, this study used the ComeJisyo medical dictionary, which, since November 2018, has 75,861 registered words. Nevertheless, depending on the domain and task, the dictionary registration of multi-word terms is inadequate, as found in this study.

Third, the Word2Vec technology presents a problem. Because Word2Vec uses random numbers during training, minute differences may occur each time the training is conducted, and the reproducibility of the study cannot be adequately guaranteed. In addition, because we did not mention the differences in research results that are due to differences in parameters, it cannot be asserted that the parameter settings used in this study are optimal.

Fourth, the ICD-10 classification prepared as an external validity measure is sometimes inappropriate for creating a clinical differential diagnosis list/cluster. Conversely, the true pre-prepared clusters mentioned in the previous study are not always found in textbooks or international classifications. If a physicians’ definitive differential diagnoses list exists, it may be worth examining external validity scales on that list.

## 5 Conclusion

The word embedded vectors produced by Word2Vec, trained on the infarction domain from a Japanese academic medical corpus, allowed us to represent the objective similarity between diseases (inter-disease distance) that can be used in PCS. The internal validity was maximized when the metric was euclidean and the method was centroid. AMI as the external validity scale with ICD-10 was maximized when the metric was cosine and correlation, and the method was average and weighted. Both NMI and ARI were maximized when the metric was cosine and the method was complete. When the frequency of word occurrence is considered, the representation of the inter-disease distance by Euclidean distance, which does not ignore the L2 norm of the embedding vector, may be a quantitative index in implementing PCS. When clinical differences similar to ICD-10 are considered, the inter-disease distance expression may be that of using the cosine distance.

## Acknowledgments

None

## Contributorship Statement

DY was responsible for the design and architecture of the study. DY undertook data collection. DY, YY, KN, TU, YO, KS, TT, and MI integrated and interpreted the results. DY and MI wrote the manuscript. DY, YY, KN, TU, YO, KS, TT, and MI read and approved the final manuscript.

## Data Availability Statement

The data used in the analysis can be downloaded from Ichushi Web, but the Japan Medical Abstracts Society (Ichushi) owns the rights to the data and cannot provide them through the authors. Embedded vectors of the training results may be provided by the authors if the Japan Medical Abstracts Society permits. The scripts used in the analysis can be provided by the authors upon reasonable request.

